# ApoDock: Ligand-Conditioned Sidechain Packing for Flexible Molecular Docking

**DOI:** 10.1101/2024.11.22.624942

**Authors:** Ding Luo, Xiaoyang Qu, Dexin Lu, Yiqiu Wang, Lina Dong, Binju Wang

## Abstract

Molecular docking is a crucial technique for elucidating protein-ligand interactions. Machine learning-based docking methods offer promising advantages over traditional approaches, with significant potential for further development. However, many current machine learning-based methods face challenges in ensuring the physical plausibility of generated docking poses. Additionally, accommodating protein flexibility remains difficult for existing methods, limiting their effectiveness in real-world scenarios. Diffusion based models has already show a good solution of those problems, such as Alphafold3(AF3). Herein, we present ApoDock, a modular docking paradigm that combines machine learning-driven conditional sidechain packing based on protein backbone and ligand information with traditional sampling methods to ensure physically realistic poses. With accurate sidechain packing and physical based pose sampling, ApoDock demonstrates competitive performance across diverse applications, highlighting its potential as a valuable tool for protein-ligand binding studies and related applications.

## Introduction

Understanding protein–small molecule interaction is crucial for comprehending biological functions such as enzyme catalysis^1,2^, signal transduction^3^, the evaluation or screening of potential new drugs^4,5^, and the design of new drugs^6^ or proteins^7,8^. Experimental methods like X-ray diffraction and cryogenic electron microscopy (cryo-EM) can reveal detailed protein–ligand interactions with atomic-level accuracy^9^, but they are expensive and time-consuming. Molecular docking is an important in silico method to rapidly model protein–ligand complexes and evaluate their binding affinities. Over the past several decades, docking has become the most common approach for modeling protein–ligand complexes^10–12^.

Classical docking methods mainly consist of two parts: sampling and scoring. The sampling component allows for ligand translation, rotation, and bond torsion within the binding pocket. The scoring component uses scoring functions (SFs) and optimization algorithms to guide the sampling process, gradually optimizing the binding poses. Recently, the emergence of machine learning-based docking methods ^13^ enables the docking pose prediction with high efficiency, which is promising for large virtual screening tasks^14^. However, when the binding site is known, the deep learning docking methods have not yet outperformed classical methods.^15–17^ In addition, it was found that no deep learning-based method currently outperforms classical docking tools in terms of physical plausibility of docked poses, as DP models were trained without knowledge of physical constraints. On the contrary, the traditional docking methods utilize physics-based functions, thus generating poses with higher physical validity.

Despite their physical plausibility, the traditional docking protocols normally treat proteins as rigid, which can limit their docking performance, especially for systems that the ligand binding may induce significant reorganization of the target protein pocket. In terms of deep learning (DL)-based flexible docking methods, the binding pose is usually reconstructed by modeling pairwise information between Coarse-graining residues and ligand atoms.^18,19^ However, such method may lead to significant clashes between ligands and residue sidechains, thereby leading to the unphysical binding pose.^20^AlphaFold2(AF2)^21^ based methods represent residues and ligand atoms as free-floating rigid bodies and predict their relative rotations and translations to construct docking poses without requiring any structural information,^15,22^ which can be taken as flexible docking process. But these methods do not surpass traditional docking methods like Vina on the PoseBuster test set.^20^ The diffusion based models also show promising results for modelling protein-ligand binding pose.^23–27^ The recently released AlphaFold3 (AF3) has taken an important step for prediction of biomolecular interactions.^28^ It can predict complexes of almost all types of molecules in the PDB database with high accuracy. Additionally, it replaces the AF2 structural module, which operates on amino acid-specific frames and side-chain torsion angles with a conditional diffusion module that directly predicts raw atomic coordinates.^29^ AF3 demonstrated significant advantages on the PoseBuster test set. However, because its code is not open source, its generalization performance on tasks such as virtual screening and binding affinity prediction has not yet known.

In most cases, the ligand binding to proteins can induce the conformational changes of the sidechains changes, while the main chains are relatively rigid.^30^ Therefore, if we can reliably model the side-chain conformers of the holo structure in the presence of ligand, the performance of semi-flexible docking methods can be further improved. However, existing side-chain packing methods were not designed for docking purposes, as the ligand binding information was not included.^31–33^ To this end, we propose a flexible docking protocol called ApoDock. ApoDock consists of three stages (Fig. 1a). 1) we introduce a conditional side-chain packing model named ApoPack, based on a Message Passing Neural Network (MPNN). ApoPack predicts the holo side-chain conformers using protein pocket backbone information and 2D ligand information. 2) we utilize the excellent re-docking capabilities of classical docking software to generate candidate poses within the predicted holo pockets. 3) we develop a Mixture Density Network (MDN)-based scoring function, ApoScore, to re-rank the generated candidate poses. This comprehensive process ensures the flexibility of the sidechains, the physical validity of the poses, and the accuracy of the results. ApoDock has demonstrated excellent docking capabilities in various application scenarios.

**Fig. 1.**
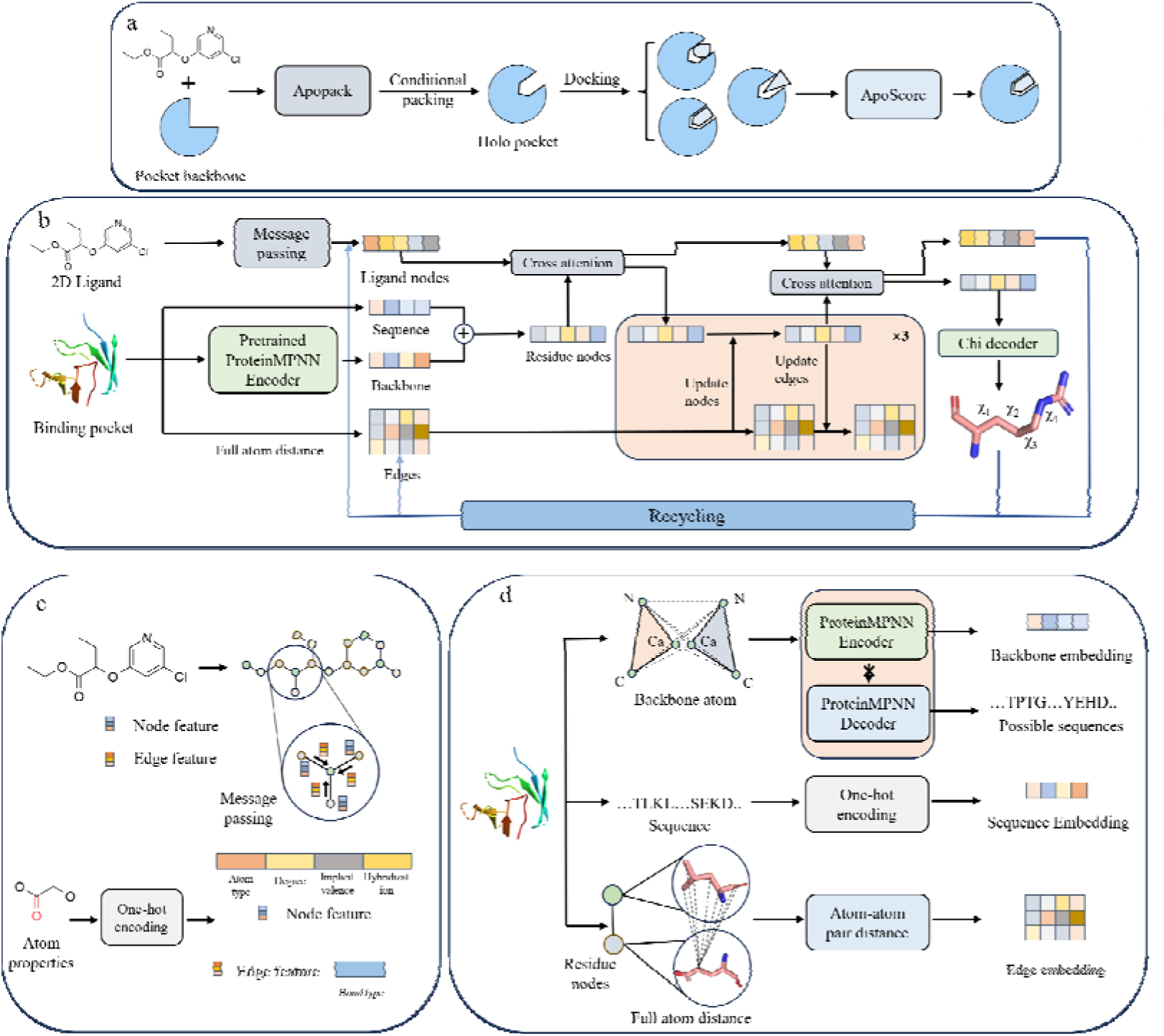
The architecture and input features of ApoDock. (a) Overview of ApoDock for flexible docking. The whole model inputs a 2d ligand and pocket backbone, and output candidate binding poses. (b) The ApockPack update the ligand node and protein node representations through MPNN layer and predict the sidechain torsion angles. (c) The ApockPack embed ligand features by atom properties and update ligand features through MPNN layer. (d) The ApockPack use backbone, sequences, and atom distances between residues as input features.

## Results

### Model architecture

ApoDock consists of two deep learning modules—ApoPack and ApoScore, and a conformation sampling module, usually Gnina^17^ or Smina^34^ (Fig. 1a). We use ApoPack to account for conformational changes when different ligands were docked into the pocket. The packed pockets are then clustered based on the root mean square deviation (RMSD) values of the sidechains. Next, we use all-atom-based physical docking software to initially generate candidate docking poses on the packed pockets. Finally, we use ApoScore to rescore and re-rank the generated poses. ApoScore is a mixture density network (MDN)-based scoring function; details of ApoScore can be found in the Supplementary Information (SI) section S3.

ApoPack is a conditional side-chain packing module that outputs different side-chain conformations based on 2D ligand information and pocket backbone information. As depicted in Fig. 1b, ApoPack embeds and updates the ligand nodes and pocket residue-level nodes using the graph neural network. Ligand nodes are first updated by a message passing neural network (MPNN). The ligand nodes and pocket nodes then interact through a cross-attention layer. The updated information is further processed by the MPNN layer, followed by a cross-attention step to iteratively update the ligand and pocket nodes (see Methods). We use a multilayer perceptron (MLP) layer to predict four torsion angles χ of every side chain. Following AlphaFold2, the output ligand nodes and predicted side-chain atomic distances are recycled as input to the model. We backpropagate the model parameters using loss functions after three cycles. Ligand encoding utilizes atom properties and atom types as node and edge features. To better represent the pocket backbone information, we use a pretrained ProteinMPNN^35^ encoder to represent pocket backbone features. ProteinMPNN embeds the protein backbone information through an encoder layer and autoregressively predicts possible sequences via a decoder layer in a determined backbone (Fig. 1d), thus it has good representation capabilities for the protein backbone.

### Packing performance of ApoPack and Rescoring Performance of ApoScore

To determine the appropriate sample number for ApoPack in docking tasks, we evaluated the relationship between the sample number and side-chain accuracy on the PDBbind time-split test set. As shown in Fig. 2a, the success rate of side-chain predictions gradually increases until the sampling number reaches 40. Therefore, we decided to sample 40 side-chain conformations of the pocket in the docking pipeline for clustering. Compared to the physically guided scoring function method Rosetta FastRelax,^36^ ApoPack shows significantly higher side-chain accuracy (Fig. 2b) and lower RMSD values (Fig. 2c), demonstrating its superior performance. The ablation study of ApoPack in Table S1 show that the importance of the cross-attention module, recycling module, and pretrained ProteinMPNN features. All of those modules are important for the sidechain packing accuracy, especially the ligand information obtained from cross-attention module, and pretrained ProteinMPNN features. Additionally, ApoPack is several hundred times faster than Rosetta FastRelax. These results demonstrate that ApoPack can predict the sidechains of protein pockets more accurately and efficiently, which can be utilized in other related tasks.

**Fig. 2:**
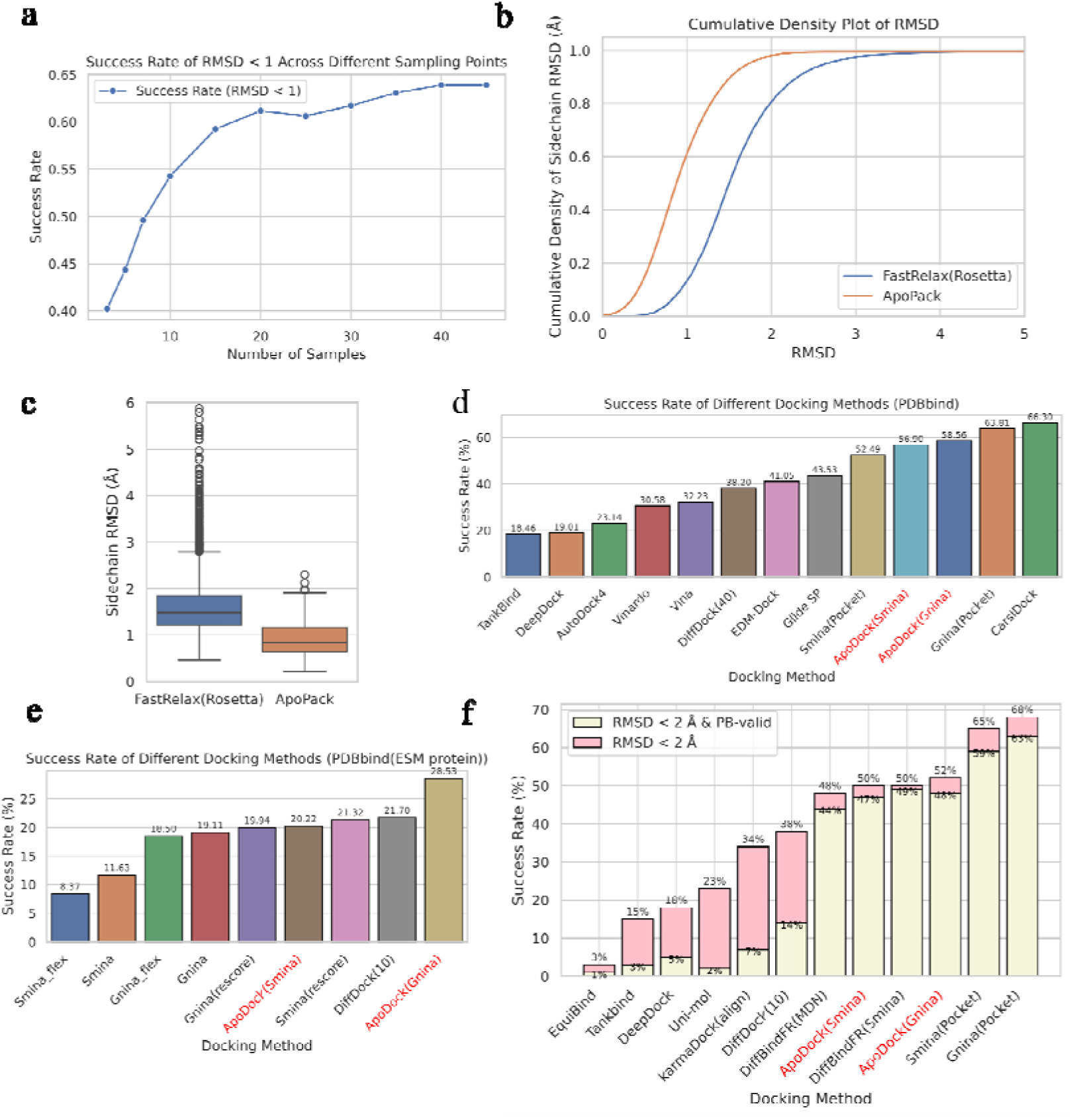
Performance of ApoDock in PDBbind and Posebusters test set. (a) Success rate (RMSD < 1 Å of all sidechain) of sidechain packing results using different sampling numbers; (b) RMSD cumulative distribution plots of Rosetta FastRelax and ApoPack; (c) RMSD distribution box plots of Rosetta FastRelax and ApoPack; Docking accuracy (RMSD < 2 Å of ligand RMSD) of different methods in PDBBind time split test set using corresponding holo structure (d) or ESMFold predicted structure (e); Except for Gnina, Smina, and ApoDock, the success rate of other methods is copy form CarsiDock. (f) Redocking performance in Posebuster test set.

The rescoring module, ApoScore, was developed to select the correct pose from the initial docking results; it determines which pose should rank at the top. Therefore, it is important to evaluate its docking power. We tested ApoScore on the most commonly used benchmark, CASF-2016.^37^ The results of docking, screening, and reverse screening power can be found in Fig. S2. ApoScore shows the best docking power with a top1 success rate of 95.1% when crystal structures are not included for comparisons (Fig. S2a). The screening power success rate of ApoScore is 64.9% (Fig. S2b), which is comparable to RTMscore (66.7%).^38^ Considering the Enrichment Factor (EF), ApoScore exhibits the best EF 1% of 29.15 (Fig. S2c). ApoScore also shows superior reverse screening power, with a top-1 success rate of 42.8%. These results demonstrate the superior performance of ApoScore. Therefore, ApoScore effectively fulfills its role of ranking poses in ApoDock.

### Redocking Performance in PDBbind and Posebuster test set

We first evaluated the docking performance of our method on the PDBbind time-split test set. We compared our method with traditional docking methods and deep learning methods that either generate poses directly or sample docking poses guided by traditional or deep learning-based scoring functions. The docking settings of those baseline model can be found in SI section S1. Fig. 2d shows the success rates of different methods. ApoDock achieves success rates of 56.90% (ApoDock(Smina)) and 58.56% (ApoDock(Gnina)), respectively, which are only lower than the redocking results of CarsiDock^39^ and Gnina.^17^ It is worth noting that the results achieved by ApoDock are obtained without prior knowledge of side-chain conformations, as ApoDock predicts side-chain conformations from ligand and pocket backbone information rather than using the provided holo structure. Therefore, except for ApoDock, the docking results of other methods in Fig. 2d can be considered as the upper limit of docking capability, given that the holo protein structures of docked poses are not available in realistic situations.

Considering a more realistic application scenario, we also compared the docking success rates of several related docking methods (Smina, Gnina, DiffDock^40^) with ApoDock on ESMFold modeled protein structures.^41^ As shown in Figure 2e, all models exhibit a lower success rate due to the limited accuracy of the modeled protein structures. Fig. S4 demonstrates that some of the ESMFold modeled protein structures have large backbone RMSDs compared to the crystal structures, and the side-chain differences are even more pronounced. Among the methods, ApoDock(Gnina) shows the best docking performance with a success rate of 28.53%. In contrast, the semi-flexible docking settings of Gnina and Smina, as well as DiffDock, achieve lower success rates because they do not consider the side-chain differences between modeled structures and crystal holo structures. We also find that ApoScore can improve the docking success rate of Smina but is limited in Gnina. This is because the scoring function of Smina is not as powerful as that of Gnina. ApoScore can rescue some poses sampled by Smina, demonstrating the superiority of ApoScore. In summary, ApoDock shows strong docking capabilities when using either holo structures or modeled structures, proving its versatility with different receptors for docking.

To better compare our method with other docking approaches, we tested ApoDock on the PoseBuster test set.^20^ A successful docking result on this test set requires not only a ligand RMSD of less than 2□Å but also compliance with steric and energetic criteria: the ligand conformation should be physically validated and not clash with the protein. As shown in Fig.2f, ApoDock achieves a top-1 docking success rate of 50% (ApoDock(Smina)) and 52% (ApoDock(Gnina)), which are higher than those of other deep learning methods. When considering pose validity, the success rates of ApoDock slightly decrease to 47% and 48%. This contrasts with other deep learning methods that show high ratios of RMSD□<□2□Å but low pose validity ratios.^20^ This may be attributed to the use of a physics-based sampling module in our method, which improves the physical validity of the generated poses.

Since ApoDock can predict the correct pocket sidechains first, followed by docking and rescoring, it tackles a more difficult task compared to other methods that utilize the crystal holo structure as the receptor for docking. Because Gnina uses the holo protein structure for redocking, it achieves a top-1 success rate (RMSD□<□2□Å) of 68%, when consider the PoseBuster validity (PB valid), the success rate decreases to 63%, which is higher than other docking methods. We also tested the use of the ApoScore module to re-rank the docked poses generated by Smina or Gnina and found that Smina + ApoScore and Gnina + ApoScore achieve docking success rates (RMSD□<□2□Å□/ PB valid) of 69%/63% and 75%/67%, respectively. The increased success rate indicates the superior rescoring power of ApoScore. The competitive results achieved by ApoDock demonstrate that it can predict side-chain conformations and rank sampled poses correctly with good physical validity.

### Performance in Apo2Holo test set

We tested ApoDock and compared it to other deep learning and traditional docking methods on the Apo2Holo test set.^42^ Unlike the redocking test set, this test set utilizes the apo structures as receptors to dock and align the generated docking poses with the holo crystal ligands to assess docking performance. Since the holo protein structures for ligand binding do not exist in real docking tasks, this test set more realistically reflects the docking capabilities of different methods. The Apo2Holo test set was divided into three groups based on the Cα RMSD of the proteins.

As shown in Fig.3a and b, ApoDock significantly reduces the mean RMSD and improves the success rate in Group 1 and Group 2 when using either Smina or Gnina as the conformation sampling method. We also tested the contributions of ApoPack and ApoScore within ApoDock. We found that the model’s performance decreased when either ApoPack or ApoScore was omitted. ApoPack with Smina seems to have lower performance compared to ApoPack with Gnina, as Gnina uses CNN scoring functions in the final pose rescoring and clustering. Compared to the traditional scoring method Smina, Gnina exhibits better performance in cross-docking test sets.^17^ We observed that using ApoScore to rescore the poses docked by Gnina or Smina also showed good performance, as the MDN-based scoring functions are competitive in cross-docking test sets.^38,39^ The significant contributions of both ApoPack and ApoScore demonstrate the superiority of ApoDock in more realistic docking tasks. The performance of ApoDock in Group 3 is not significantly better than other methods in Fig. 3 and Table 1, as the Cα RMSD between the apo and holo structures is larger (>1.5□Å). Our model cannot account for the flexibility of the main chain, making it difficult to obtain correct poses in some cases.

**Fig. 3.**
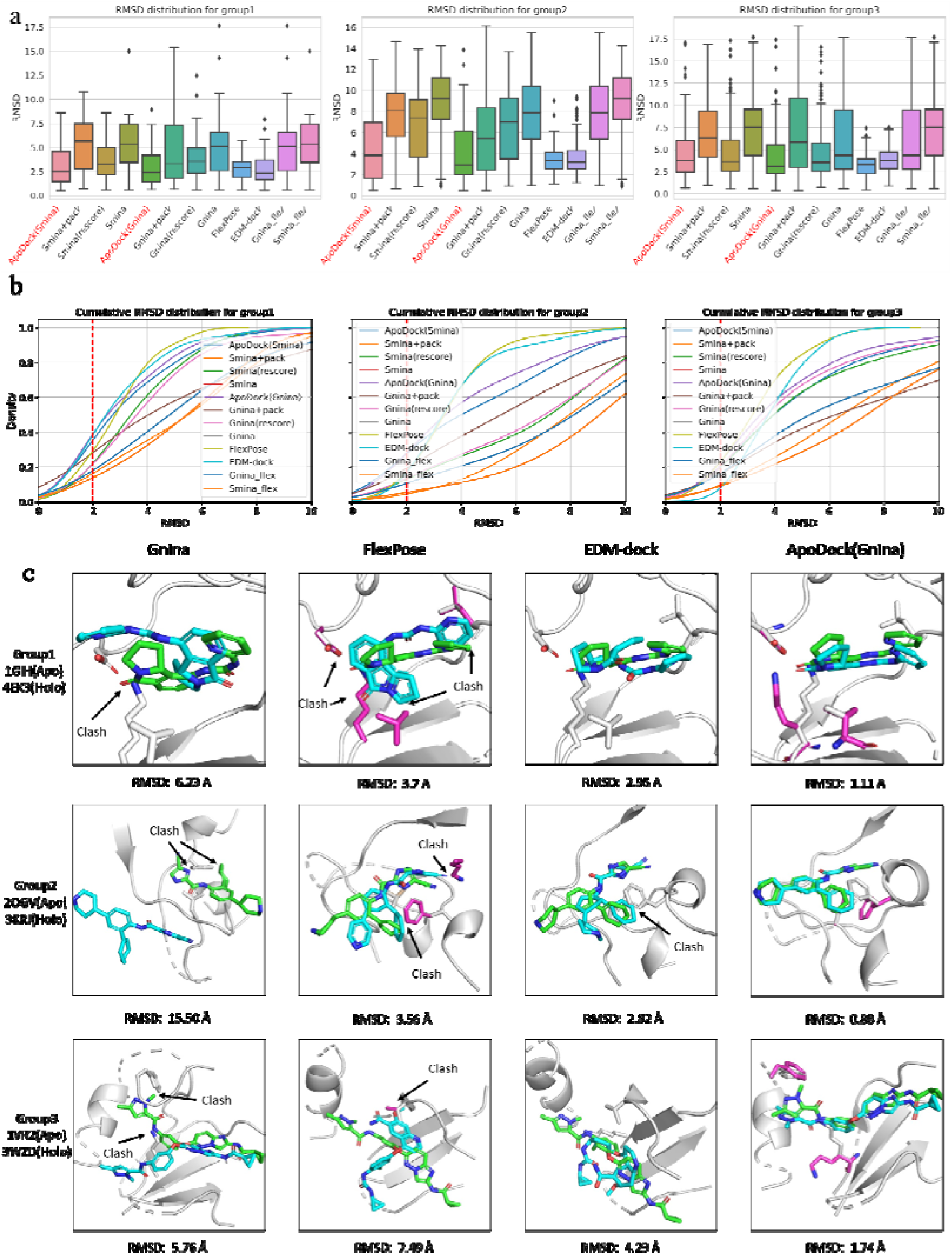
Prediction accuracy of different docking method in Apo2holo test set. RMSD distribution box plots (a) and cumulative distribution plots (b) of different docking methods on three groups; (c) Examples of docking poses in three groups obtained by representative docking method. Green represents crystal pose, cyan represents docked top1 pose, purple represents the packed sidechain by the flexible docking method, the gray represents the apo structure.

**Table 1:** Performance comparison of different method in Apo2Holo test set.

| Method | Group1 |  |  |  | Group2 |  |  |  | Group3 |  |  |  |
| --- | --- | --- | --- | --- | --- | --- | --- | --- | --- | --- | --- | --- |
|  | Top1<br>RMSD |  | Top5<br>RMSD |  | Top1 RMSD |  | Top5 RMSD |  | Top1 RMSD |  | Top5 RMSD |  |
|  | %<2 | Med | %<2 | Med | %<2 | Med | %<2 | Med | %<2 | Med | %<2 | Med |
| Smina+Pack | 11.1 | 5.32 | 20.3 | 4.27 | 3.60 | 7.97 | 18.32 | 5.75 | 8.28 | 6.71 | 14.1 | 5.05 |
| Smina+rescore | 22.2 | 3.61 | 27.8 | 3.04 | 14.6 | 6.68 | 17.17 | 5.88 | 15.90 | 4.79 | 24.82 | 4.11 |
| Smina | 13.0 | 5.33 | 24.1 | 3.87 | 6.00 | 8.61 | 9.12 | 6.73 | 8.28 | 7.30 | 14.64 | 5.30 |
| Gnina+Pack | 31.2 | 4.86 | 44.4 | 3.17 | 18.3 | 5.81 | 26.8 | 4.38 | 10.2 | 7.20 | 19.84 | 5.31 |
| Gnina +rescore | 22.2 | 3.89 | 29.6 | 3.17 | 14.6 | 6.60 | 20.7 | 5.59 | 14.6 | 4.52 | 24.2 | 4.03 |
| Gnina | 14.8 | 5.21 | 29.6 | 3.56 | 13.4 | 7.67 | 17.1 | 6.51 | 10.2 | 6.47 | 18.50 | 4.82 |
| Gnina_flex | 23.9 | 5.42 | 32.6 | 3.13 | 13.5 | 6.28 | 16.2 | 5.26 | 13.4 | 6.78 | 21.0 | 4.90 |
| Smina_flex | 16.7 | 4.98 | 31.5 | 3.20 | 6.10 | 7.38 | 15.85 | 5.29 | 14.65 | 6.1 | 22.29 | 4.41 |
| FlexPose | 18.5 | 2.87 | - | - | **7.3** | **3.53** | - | - | 11.1 | **3.26** |  | - |
| EDM-dock | 38.8 | **2.81** | - | - | 13.4 | 3.66 | - | - | 4.4 | 3.84 | - | - |
| Apodock(Smina) | 38.9 | 3.14 | 50.0 | 2.53 | **31.7** | 4.47 | 36.5 | 3.66 | 16.6 | 4.64 | 24.8 | 3.75 |
| Apodock(Gnina) | **42.6** | 2.94 | **51.9** | **2.33** | 23.2 | **4.17** | **37.8** | **3.50** | **19.7** | 4.16 | **29.9** | **3.58** |

Deep learning docking methods that do not have sampling stage, such as FlexPose^43^ and EDM-dock,^19^ have lower mean RMSDs compared to other methods (Fig. 3 and Table 1). However, lot of poses generated by FlexPose could not be processed for docking or RMSD calculation due to incorrect bond connections and bond lengths. Implausible poses can also be found in EDM-dock (Fig. 3c). While deep learning models seem to generate poses with lower mean RMSDs overall, they have a lower success rate when considering ligand RMSD < 2□Å (Table 1). This is because these machine learning methods were trained to minimize topological information or atomic distances between the predicted ligand and the crystal ligand but ignored physical constraints in both intermolecular and intramolecular interactions. Three examples can be found in Fig. 3c for each group.

ApoDock generates poses with lower RMSDs and improved pose plausibility (Fig. 3), because it recovers the conformations of important sidechains, utilizes physically constrained traditional docking methods for sampling, and employs state-of-the-art scoring functions to rescore docked poses.

### Performance in crossdocking test

Crossdocking tasks represent another realistic docking scenario. In cross-docking, a ligand is docked to its cognate receptor, but the side-chain conformation of the receptor may differ from that in the ideal (holo) conformation. Herein, we compare ApoDock with Gnina and Smina and analyze the contributions of the modules within ApoDock. As shown in Fig. 4a and b, ApoDock (Gnina/Smina) achieves top-1 success rates of 46.6% and 46.0%, respectively, with lower mean RMSD values compared to other methods. Notably, Gnina without packing or rescoring achieves a success rate of 38.8%, which is higher than that of the traditional docking method Smina (27.7%). Gnina was trained on the CrossDocked2020 dataset because this dataset enhances and expands the available data by including ligand poses cross-docked against noncognate receptor structures and by purposefully generating counterexample poses. These features help the model better learn to distinguish correct from incorrect docking patterns. Therefore, it performs well in cross-docking tasks. We also find that rescoring the poses generated by Gnina or Smina using ApoScore shows significant improvement (from 38.8% to 45.2% for Gnina, and from 27.7% to 43.1% for Smina). This proves that the scoring function constrains the docking power of the two methods.

**Fig. 4.**
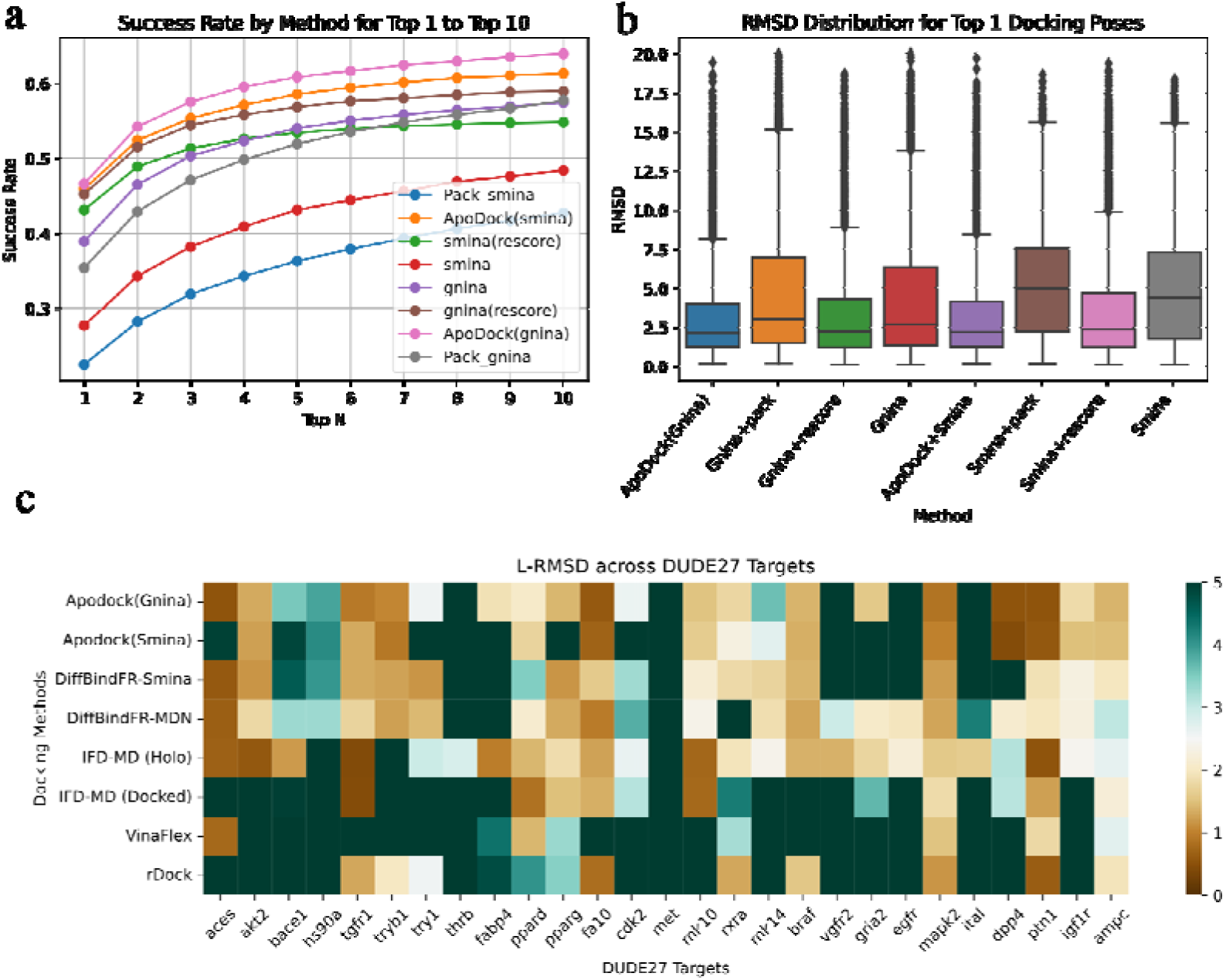
(a) Success rate and (b) top1 RMSD cumulative distribution of Smina, Gnina, ApoDock, and module ablactations in Crossdocking test set. topN is the percentage of targets ranked above or at N with a RMSD less than 2 Å. (c): RMSD values of 27 DUDE-AF2 targets by different docking methods.

Unlike what was observed in the Apo2Holo test set, the ApoPack module does not seem to have a significant improvement on the docking results of Gnina/Smina with rescoring (from 45.2% to 46.6% for Gnina, and from 43.1% to 46.0% for Smina), as the difference in side-chain conformation between the holo structures in the cross-docking test set is not as significant as in the Apo2Holo test set. Without ApoScore, ApoPack combined with Gnina/Smina has a relatively lower success rate compared to Gnina/Smina alone. This is due to the fact that predicting side-chain conformations is not as accurate as using holo side-chain conformations from homologous proteins in some cases. However, by combining ApoPack, Smina/Gnina, and ApoScore (i.e., ApoDock), good docking results can be achieved on both apo and holo structures. Moreover, holo structures are not always available. The good performance of ApoDock in the cross-docking test set demonstrates its practicality in real docking scenarios.

### Performance in AF2-DUDE test set

To further evaluate the docking performance on AF2-modeled proteins, we compared our model with several flexible docking methods on the DUDE-AF2 test set. Zhang et al. used this test set to assess the induced-fit-based protocol (IFD-MD) method and found that refining the AF2-modeled protein can improve docking accuracy close to that of the holo structure.^44^ IFD-MD is a time-consuming method that requires long MD simulations and also needs ligand pose templates to induce the protein conformation.^45^ In Fig. 4c, IFD-MD shows good performance when using the crystal ligand as the template, but the crystal ligand binding pose is not available in advance in real application scenarios. A more realistic result for IFD-MD is to use the docking pose as the template for IFD-MD (referred to as IFD-MD (docked)). It was found in Fig. 4c that the docking performance of IFD-MD significantly decreased when using the docking pose as the template. This proves that IFD-MD performance seriously depends on the initial pose. As can be seen in Fig. 4c, ApoDock shows comparable performance with DiffBindFR^23^ and IFD-MD (docked). Compared to DiffBindFR-MDN, ApoDock (Gnina) shows lower RMSD values in the FABP4 and RXRA targets. This proves that our model can be complementary to DiffBindFR, and the relevant results need to be further explored. We analyzed some targets where IFD-MD succeeded (RMSD < 2□Å) but ApoDock (Gnina) failed, and we found that most of the crystal ligands have obvious clashes with the protein main chain of AF2-modeled proteins in these targets (Fig. S5). Therefore, it is impossible to sample a correct pose with physical validity. In summary, these results suggest that our model has superior docking ability when considering AF2-modeled proteins as receptors.

### Docking time comparison

The total docking time of our method is about several times that of semi-flexible docking methods. As seen in Fig. S3a, compared to other flexible docking methods, our method has acceptable docking times. FlexPose and EDM-dock are very fast, as they generate poses directly without sampling. The docking speed of our method falls between that of deep learning methods and traditional flexible sampling methods, suggesting it offers a good balance between docking time and pose validity.

ApoDock (Gnina) has a longer docking time compared to ApoDock (Smina) because Gnina performs computations with single (32-bit) precision rather than double (64-bit) precision due to the need to shift calculations to the GPU for efficient CNN scoring. As shown in Fig. S3b, most of the time spent by ApoDock (Gnina) is on the sampling part. Our modules, ApoPack and ApoScore, account for only 4.5% and 0.6% of the total docking time, respectively. The docking time is also dependent on the number of samples. Thus, the main limitation of our method is the sampling part; employing faster sampling methods would improve the docking efficiency of ApoDock.

## Discussion

We developed ApoDock, a flexible docking method that can generate docking poses with accuracy and physical plausibility. ApoDock demonstrates impressive docking power in redocking, docking with apo structures, cross-docking, and model-docking tasks. The satisfactory performance of ApoDock is due to the correct prediction of side-chain conformations by the developed ApoPack module, aided by state-of-the-art sampling software and the SF ApoScore. ApoPack borrows many ideas from AF2 in its architecture, such as the recycling mechanism and predicting torsion angles of sidechains. In terms of input features, the pre-training features provided by ProteinMPNN are also crucial for the performance of ApoPack.

The redocking test results show that current deep learning algorithms do not completely surpass traditional algorithms, especially when the conformation of the protein pocket are known; traditional docking methods like Gnina remain competitive. ApoDock exhibits slightly decreased performance compared to Gnina or Smina when using holo structures for docking (Fig. 2d). However, using ApoScore can significantly improve the redocking results of Gnina or Smina (Fig. 2e, 3a, and 3b), indicating that the scoring function limits the docking performance of traditional methods.

Performance on the Apo2Holo test set demonstrates that both the ApoPack and ApoDock modules play essential roles in enhancing the docking success rate. While the ApoScore module contributes to increased success rates on the Apo2Holo test set, the inclusion of the ApoPack module significantly raises the proportion of correct poses, leading to an overall improvement in docking outcomes (Fig. 3b). In comparison to Gnina, which performs exceptionally well in crossdocking, ApoDock also shows competitive performance. However, the advantages of ApoPack in crossdocking scenarios are less pronounced than in docking tasks involving apo structures or AF2-modeled structures. This difference may arise from the minimal structural deviations between holo structures of alternative ligands and the target holo structure. In those case, the traditional semi-flex docking method can show good performance as well.

The time spent on ApoDock docking mainly arises from the sampling time of traditional docking software, which depends on the number of pocket clusters. Our approach has moderate docking times compared to traditional docking methods and deep learning techniques. While deep learning docking methods are very fast, they can easily overlook physical validity. Incorporating physical constraints into the pose generation process guided by deep learning algorithms is an important direction for future improvements.

Although AF3 has shown overwhelming results in protein–ligand docking tasks, its screening power remains unknown. The confidence scores in AF3 may correlate with screening ability. However, the reliance on multiple sequence alignments (MSA) and the slower modeling process may limit the application of AF3 in virtual screening scenarios. Leveraging ApoDock’s strong virtual screening and docking capabilities, our model can serve as a supplement to AF3. In summary, our method provides a powerful tool for addressing protein–ligand binding-related problems.

## Methods

### Data sets

#### Training set

We used PDBbind v2020 to train our packing model and rescoring model.^46^ Following EquiBind’s training strategy,^47^ we used 363 complex structures uploaded after 2019 as the test set, excluding molecules that could not be read by RDKit. The remaining complexes were divided into a training set and a validation set. The training set included 16,334 complexes, and the validation set included 964 complexes. We refer to this dataset as the "PDBbind time-split dataset." To better benchmark against other scoring functions, we also trained a version of the MDN scoring function using the PDBbind core set as the test set.

#### PoseBuster Test Set

The PoseBuster dataset comprises publicly available crystallographic complexes carefully selected from the PDB database.^20^ It includes only complexes published since 2021, ensuring that it does not contain any complexes present in the PDBbind General Set v2020 used for training in many methods. The dataset undergoes a series of standard quality checks using the well-established cheminformatics toolkit RDKit.^48^ The PoseBuster test suite verifies the chemical and geometric consistency of the ligands—including their stereochemistry—and assesses the physical plausibility of intra- and intermolecular interactions.

#### Apo2Holo Test Set

Zhang et al. collected 32 pairs of apo and holo structures for the DUD-E dataset from the PDB database.^42^ The structures in this dataset are divided into three categories based on the Cα RMSD: Group 1 (Cα RMSD ≤ 0.5□Å), Group 2 (0.5□Å < Cα RMSD ≤ 1.5□Å), and Group 3 (Cα RMSD > 1.5□Å). This dataset effectively simulates the use of apo structures for docking in virtual screening situations when the holo structure is unknown.

#### Cross-Docking Test Set

The cross-docking dataset recently published by Wierbowski et al. provides a meaningful way to evaluate real docking scenarios.^49^ Proteins with different holo structures are superimposed onto the correct complex crystal structure, and the superimposed holo proteins are used as receptors during docking. The dataset consists of 94 unique protein binding pockets and 4,399 unique ligands, with an average of 46 ligands per target. A total of 7,970 protein–ligand pairs sampled by Gnina were used to test the cross-docking performance of the model.

#### DUDE27-AF2 Test Set

The DUDE27-AF2 test set was originally used to evaluate the possibility of using IFD-MD to induce more holo structures from unliganded AF2 predictions for molecular docking.^44^ It contains a total of 27 targets. For each target, we downloaded the apo and holo structures assigned by DUD-E from the PDB and the AF2 structure from the AlphaFold Protein Structure Database.^50^ We determined the binding pocket by superimposing the AF2-predicted protein onto the correct holo structure with PyMoL^51^.

#### Data Processing

We selected the amino acid residues within 10□Å of the crystal ligand in the protein as the binding pocket for training and testing our model. For the Apo2Holo dataset, we aligned the apo and holo structures and extracted the binding pockets using PyMOL.^51^ For the protein structures predicted by ESMFold,^41^ we followed DiffDock’s alignment method.^40^ The detailed prediction parameters can be found in SI section S5.

### Input Features

We use graphs to represent both the ligand and the pocket. Ligand atoms are represented as nodes in the ligand graph, and covalent bonds are represented as edges. Pocket residues are represented as nodes in the pocket graph, and edges connect the k-nearest residues corresponding to each node.

#### Ligand Encoding

The initial ligand features for both the ApoPack and ApoScore modules consist of one-hot encodings of the atom type (C, N, O, S, F, P, Cl, Br, I, Unknown), atom degree, implicit valence, and hybridization state. The edge features are one-hot encodings of bond types.

#### Pocket Encoding

We use sequence features and backbone features as the node features for the ApoPack pocket encoding. Sequences are represented as one-hot encodings. We use the pretrained ProteinMPNN encoder to obtain the backbone features, as it is trained to generate sequences from backbone information, providing good representations of backbone structure. The edge features of the ApoPack pocket are the 14 full-atom distances between two residues. We initially set the side-chain atom distances to zero and update them through model recycling.^31^ The detailed pocket encoding of ApoScore can be found in the SI section S2.

### Model architecture

#### ApoPack architecture

We use message passing neural network (MPNN) to update ligand node features, and pocket node features. For each node *i*∈*V*, the message from its neighboring node *j*∈*N*(*i*) is computed as:

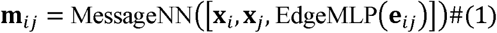

where *e_ij_* represents the edge features between nodes *i* and *j*, and [*x_j_*, *x_i_*, *e_ij_*] denotes the concatenation of the node and edge features. The transformations MessageNN and EdgeMLP are applied through MLPs. The messages from all neighbors of node i are aggregated using a summation:

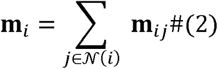

The aggregated message *m_i_* is then used to update the node features:

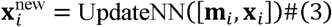

Here, the UpdateNN module applies an MLP to combine the aggregated message and the original node features. Finally, the node features are passed through an MLP for transformation, using distinct MLPs depending on the specific variant of the features:

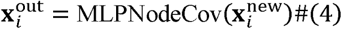

The hidden dimension is set to 128. For ligand node update, one layer update end, and every MLP linear layer will pass Dropout, LeakyReLU, and BatchNormlization layer. For pocket node update, the output Xi need to update the edge features like ProteinMPNN:

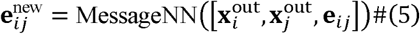

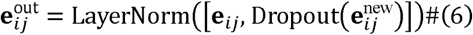

The output edge features will input to the next layer to update the node features. We select the nearest 30 residues as neighbors when update pocket nodes information. We use cross-attention block to update the ligand and pocket information:

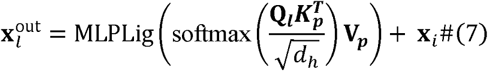

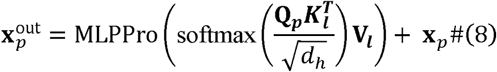

Where the *l* represents the ligand and *p* represents protein. *Q*, *K*, *V* represent the linear transform of ligand and pocket node features. The MLP linear block contains two linear, Dropout, LeakyReLU, and BatchNormlization layers.

#### ApoScore Architecture

ApoScore is a Mixture Density Network (MDN) used for scoring docked poses by utilizing 2D ligand information, pose coordinates, and 3D pocket information. The ligand is embedded in the same way as in ApoPack, and the pocket information is embedded using a GVP-GNN (Geometric Vector Perceptron Graph Neural Network) layer. Detailed implementations of ApoScore can be found in the SI section S2.

#### Loss Function of ApoPack

Inspired by AlphaFold2^21^ and PIPPack,^31^ we consider the side-chain conformation prediction task as the prediction of four *χ* torsion angles for every amino acid. We use a bin width of 5° to discretize the torsion angles from (–*π*, *π*) and predict the angle bins. Such a classification method is beneficial compared to other methods that predict side-chain conformations via dihedrals by performing a regression task on sin□*χ* and cos□*χ* for all *χ* angles. An offset value is also predicted to precisely place the angle within the bin.

### Training parameters

All the training parameters of ApoPack and ApoScore can be found in SI section S3.

## Availability of data and materials

Demo, instructions, and codes for ApoDock will be available at https://github.com/DingLuoXMU/ApoDock_public

## Authors’ contributions

D.L. and B.W. conceived and designed the experiments. D.L. performed the experiments. D.L., X.Q., D.Lu., Y.W., L.D and B.W. analyzed the data. D.L. contributed analysis tools. D.L. and B.W. wrote the paper.

## Funding and Acknowledgements

This work was supported by National Key Research and Development Program of China (2019YFA0906400) and NSFC (No. 22073077, 21933009, and 21907082).

