## Supplementary material for "ApoDock: Ligand-Conditioned Sidechain Packing for Flexible Molecular Docking": si

**Contents**

S1. Ablation study of ApoPack.

S2. Baseline models.

S3. Details of ApoScore.

S4. Training hyperparameters.

S5. Docking time evaluation.

S6. ESMFold predicted structure.

S7. Failure examples of DUDE-AF2 test set.

**S1. Ablation study of ApoPack**

Table S1: ablation study of ApoPack in PDBbind time split test set.

| Cross-attention | recycling | ProteinMPNN features | Success rate (RMSD < 1 A) | Mean RMSD |
| --- | --- | --- | --- | --- |
| √ | √ | √ | 0.63 | 0.95 |
| × | √ | √ | 0.57 | 1.24 |
| √ | × | × | 0.60 | 1.04 |
| × | × | √ | 0.54 | 1.40 |
| × | × | × | 0.50 | 1.47 |

**S2. Baseline models**

**Smina.** Smina^1^ is a fork of Autodock Vina^2^, Smina improves the empirical scoring function of Vina by fitting more structure and affinity data. It also supports input in multiple formats. Compared to Vina, Smina is faster, more accurate. Smina also supports define the docking box by a given ligand. We use “--autobox_ligand” and “--autobox_add 6” to define the docking center and box size. The input format of ligand and proteins are SDF and PDB, respectively. The num_modes was set to 40, exhaustiveness was set to 32. we repeat the docking process 10 times to get the top ranked poses.

**Gnina.** Gnina inherited the advantage of Smina.^3^ It uses CNN based scoring function to cluster and rescoring the poses in the final docking steps.^4^ The scoring function was trained in a large crossdocking dataset, so it has good docking power in different docking tasks. Gnina v1.1 was used for docking. The docking parameters was set as same as Smina.

**Gnina_flex/Smina_flex**. The flexible sidechain settings in Gnina and Smina were used for perform flexible docking. We define the flexible residues near the crystal structure of 3.5 Å. Other docking parameters was set as same as Smina.

**FlexPose.** FlexPose is an end-to-end deep learning docking method that flexible modeling of complex structures in Euclidean space without the following conventional sampling and scoring strategies.^5^ We follow the default parameters in <https://github.com/tiejundong/FlexPose>. As FlexPose were trained in mol2. format of the ligand, we convert the SDF format of ligand to mol2 format for inference. We removed the docking results in which FlexPose could not generated validity poses for calculating RMSD.

**EDM-dock.** EDM-dock is a flexible protein–ligand docking method based on the prediction of an intermolecular Euclidean distance matrix.^6^ The default setting of EMD-dock in <https://github.com/MatthewMasters/EDM-Dock> are used for docking. The box was defined as a 22.5×22.5×22.5 Å^3^ cube. Energy minimization was performed to refine the generated poses.

**Rosetta FastRelax.** Fast Relax minimizes the energy of molecular models by adjusting geometries to achieve a more stable configuration. Fast Relax allows the addition of positional or structural constraints (e.g., backbone distance or angle constraints) to maintain specific regions of the structure, preventing significant changes in critical areas. The Rosetta (version: rosetta.source.release-371) were used for packing. The command: ‘‘mpirun -np 8 relax.mpi.linuxgccrelease -database /mnt/d/code/rosetta.source.release-371/main/database -in:file:s {pdbbind_dir}/{pdbid}/{pdbid}_protein_processed.pdb -nstruct 40 -out:prefix {pdbbind_dir}/{pdbid}/{pdbid}_relax -relax:quick’’ were used.

**Other methods.** For the pdbbind redocking results, the docking results of Tankbind,^7^ DeepDock, Autodock4, Vinardo, Vina, Glide SP, and CarsiDock were reported by orgirnal Carsidock reference.^8^ The DiffDock results reported by the DiffDock reference.^9^ DiffDock (40) means samples 40 docking poses. For the posebuster docking results, Equibind, Tankbind, DeepDock, Uni-mol were reported by the posebuster reference. Except ApoDock, the results of DUDE-AF2 were reported from DiffBindFR reference.

**S3. Details of ApoScore**

Actually, ApoScore is knowledge-based scoring function, which estimate the bind pose by the statistical potential. Different from the traditional knowledge-based scoring functions, ApoScore fits the distance potential energy between the protein atom and the ligand atom through parameters learned by the neural network, rather than defined from the training data. ApoScore uses a Gaussian mixture density network (MDN) to characterize the complex distribution between protein-ligand atoms. The model only needs to learn the weight parameters, σ, μ and Pi required to fit the MDN network.

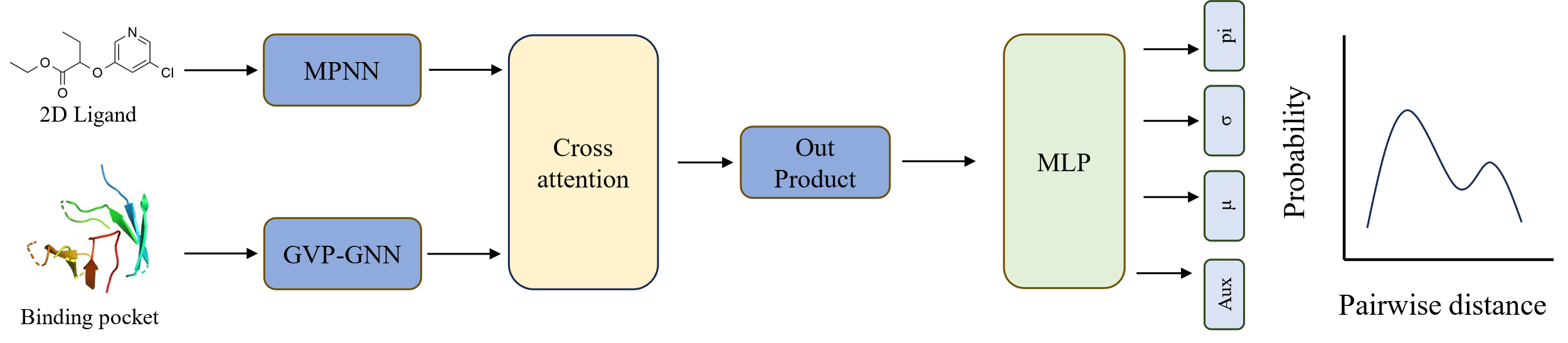

Fig. S1. Architecture of ApoScore.

The overall architecture of ApoScore is shown in Fig. S1. We use Graph Neural Network (GNN) to extract the 2D ligand and binding pocket information. The features used for MPNN and GVP-GNN can be found in Table S1. we used a cross-attention layer to strengthen the interactions between ligand features and pocket features. Then a out product operation is performed to get the interaction features. Finally, the parameters needed for MDN were obtained by a MLP layer. The statistical potential for a specific protein–ligand complex could be calculated by summing up the negative log-likelihood values of all potential protein–ligand node pairs. We adopt a similar strategy with RTMscore, which calculate the distance likelihood by the minimum distance between a specific ligand atom and each residue atom. The loss function of MDN is written as eq1:

$$\begin{aligned} \mathcal{L}_{\mathrm{MDN}}=-\log P\left( d_{p,s} | h_{p}^{\mathrm{prot}},h_{s}^{\mathrm{lig}} \right)=-\log\sum_{n=1}^{N_{s}} \rho_{p,s,n}N\left( d_{p,s} | \mu_{p,s,n},\sigma_{p,s,n} \right)\#\left( 1 \right) \end{aligned}$$

Where *p* represents protein*, s* represents ligand, *d* represents distance, *n* represent the *n-th* atom in ligand, and *h* represent the updated features.

Then, the statistical potential for a specific protein–ligand complex are calculated by summing up the negative log-likelihood values of all potential protein–ligand node pairs(eq2):

$$\begin{aligned} U_{\left( * \right)}=-\sum_{p=1}^{P} \sum_{s=1}^{S} \log P\left( d_{p,s} | h_{p}^{\mathrm{prot}},h_{s}^{\mathrm{lig}} \right)=-\mathrm{score}\#\left( 2 \right) \end{aligned}$$

We also tested the ApoScore in CASF-2016, which is a golden benchmark designed for different scoring functions. The results are shown in Fig. S2. ApoScore shows state-of-the-art (SOTA) performance compared to other deep learning methods in both docking, screening (screening power in enrichment factor), and reverse screening power. The success rate of screening power is comparable to RTMScore.^10^ Those results proves that ApoSore’s good docking ability, which can promote ApoDock to the different docking tasks, such as docking, screening, or lead compound optimization scenario.

Table S1: node and edge feature description of GNN models

| EGNN | Features | Dim^a^ | description |
| --- | --- | --- | --- |
| MPNN | **Nodes** |  |  |
|  | Atom type | 10* | Heavy atom type [C, N, O, S, F, P, Cl, Br, I, others] |
|  | Degree | 7* | Number of covalent bonds [0,1,2,3,4,5,6,] |
|  | Implicit valence | 7* | Implicit valence of the atom [0,1,2,3,4,5,6] |
|  | Hybridization | 5* | [sp, sp2, sp3, sp3d, sp3d2] |
|  | Aromatic | 1 | Whether the atom part of an aromatic system |
|  | Hydrogens | 5 | Number of connected hydrogens [0,1,2,3,4] |
|  | **Edges** |  |  |
|  | bond type | 6* | Single bond, Double Bond, Triple Bond, Aromatic, In Ring, Conjugated [0,1,2,3,4,5] |
| GVP-GNN | **Nodes** |  |  |
|  | N_scalar | 6 | [sin, cos] ◦ [φ, ψ, ω] computed from C*_i_*−1, N*_i_*, C*α_i_*, C*_i_*, and N*_i_*_+1_ |
|  | N_vector | 3 | The forward and reverse unit vectors in the directions of C*α_i_*_+1_ − C*α_i_* and C*α_i_*_−1_ − C*α_i_*, respectively;  The unit vector in the imputed direction of C*β_i_* − C*α_i_* |
|  | Sequence | 20 | A one-hot representation of amino acid identity |
|  | **Edges** |  |  |
|  | E_scalar | 32 | The encoding of the distance \|\|C*α_j_* − C*α_i_*  \|\|_2_ in terms of Gaussian radial basis functions;  A sinusoidal encoding of *j* − *i* |
|  | E_vector | 1 | The unit vector in the direction of C*α_j_* − C*α_i_* |

^a^: Dimension of feature vector.

^b^: One hot encoding of features.

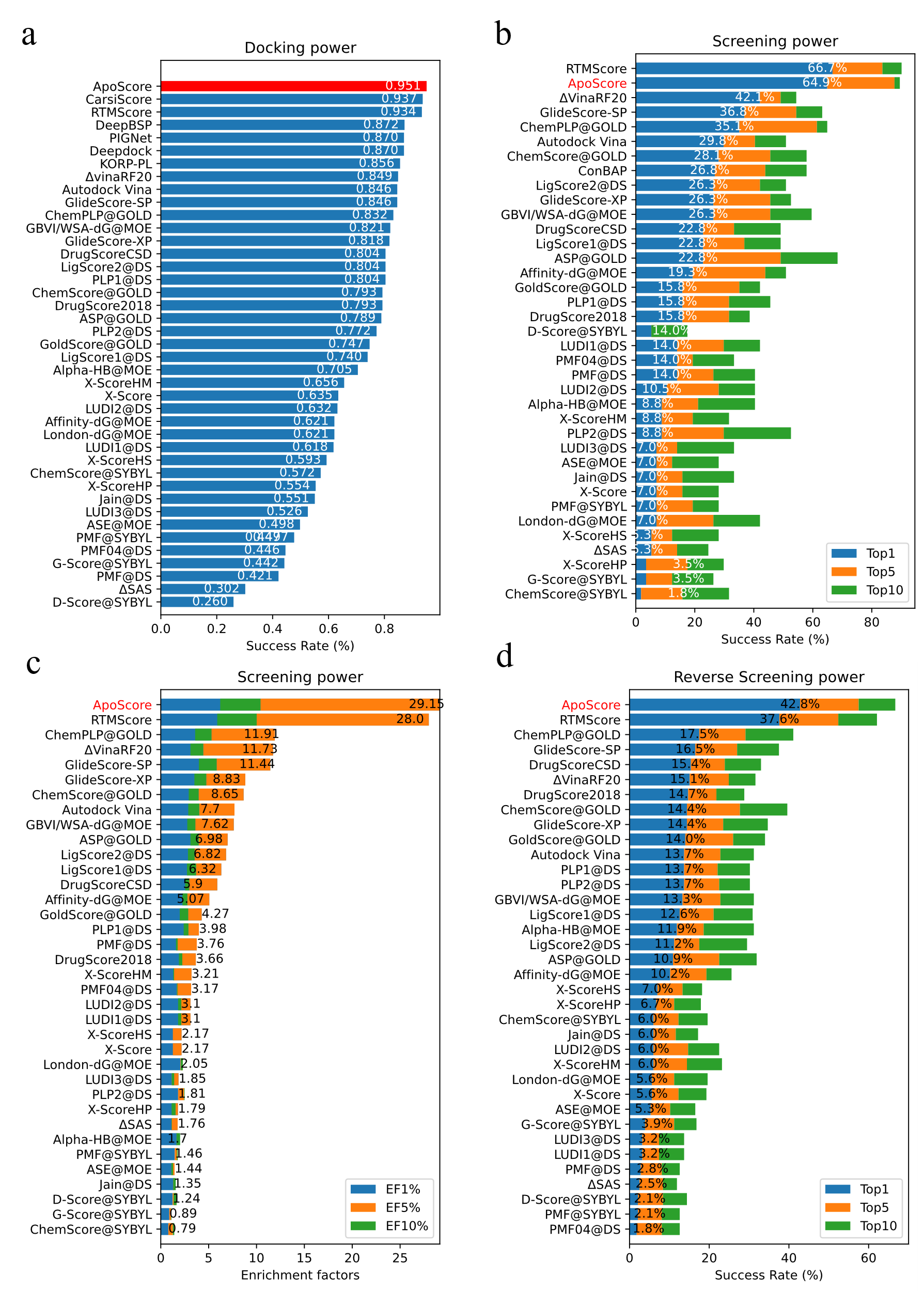

Fig. S2: The docking and screening power of ApoScore in CASF-2016 benchmark. Our method is height in right. (a) The docking power test for top1 success rate, the crystal structures are not included. (b) The success rate of the screening power. The enrichment factors (EF) of screening power. (d) The revers screening power.

**S4. Main Hyperparameters**

Table S2: Hyperparameters of ApoPack

| Model | Hyperparameters | Setting |
| --- | --- | --- |
| MPNN layer for ligand | Input dimension of node | 35 |
|  | Input dimension of edge | 6 |
|  | Hidden dimension | 256 |
|  | Number of layers | 3 |
| MPNN layer for protein | Input dimension of node | 149 |
|  | Input dimension of Edge | 3152 |
|  | Positional embeddings | 16 |
|  | Number of layers | 3 |
| Other Hyperparameters | Number of MLPs | 3 |
|  | Active function of chi decoder | RELU |
|  | Recycling time | 3 |
|  | Number of attention heads | 4 |
|  | Dropout | 0.1 |
|  | Number of chi bins | 72 |
|  | Optimizer type | Adam |
|  | Batch size | 64 |
|  | Learning rate | 0.001 |
|  | Weight decay | 0.00001 |
|  | Inference temperature | 2.0 |

Table S3: Hyperparameters of ApoScore.

| Model | Hyperparameters | Setting |
| --- | --- | --- |
| MPNN layer for ligand | Input dimension of node | 35 |
|  | Input dimension of edge | 6 |
|  | Hidden dimension | 256 |
|  | Number of layers | 3 |
| GVP-GNN for protein | Input dimension of node | 29 |
|  | Input dimension of edge | 33 |
|  | Hidden dimension | 256 |
| Other Hyperparameters | Number of gaussians | 10 |
|  | Optimizer type | Adam |
|  | Batch size | 64 |
|  | Learning rate | 0.001 |
|  | Weight decay | 0.00001 |
|  | Optimizer type | Adam |
|  | Training epochs | 2000 |
|  | Patience for early stopping | 200 |

**S5. Docking time evaluation**

The Docking time of ApoDock and other methods can be found in Fig. S3. The comparison of ApoDock and other methods are on 32 core CPU with one RTX 3090 GPU.

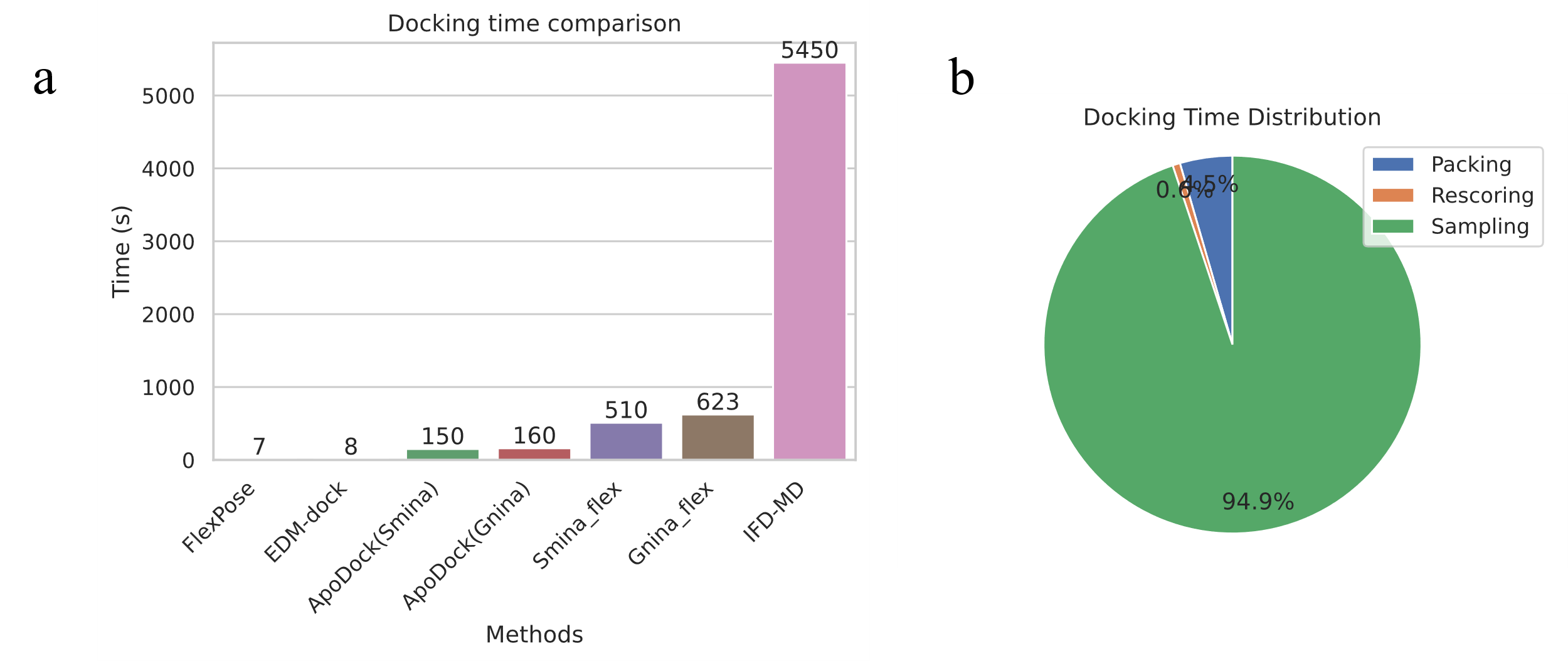

Fig.S3: Docking time evaluation of Apodock. (a) Docking time comparison of ApoDock with other methods. (b) Docking time distrubutison of ApoDock(Gnina).

**S6. ESMFold predicted structure**

We first use Biopython^11^ to extract the protein sequence of PDBs in PDBbind test set. We only consider the standard amino acid and skip the “HETATM” line. The different chains are split by “:”. We set the chunk size = 8 to predict more sequence with affordable memory as soon as possible in one RTX3090 GPU. The pLDDT distribution and Cα RMSD distribution are shown in Fig. S4. The results show that the ESMFold predicted structure is not always accurate, some of the proteins has low confidence score (pLDDT) and large Cα RMSD, which limits the docking ability.

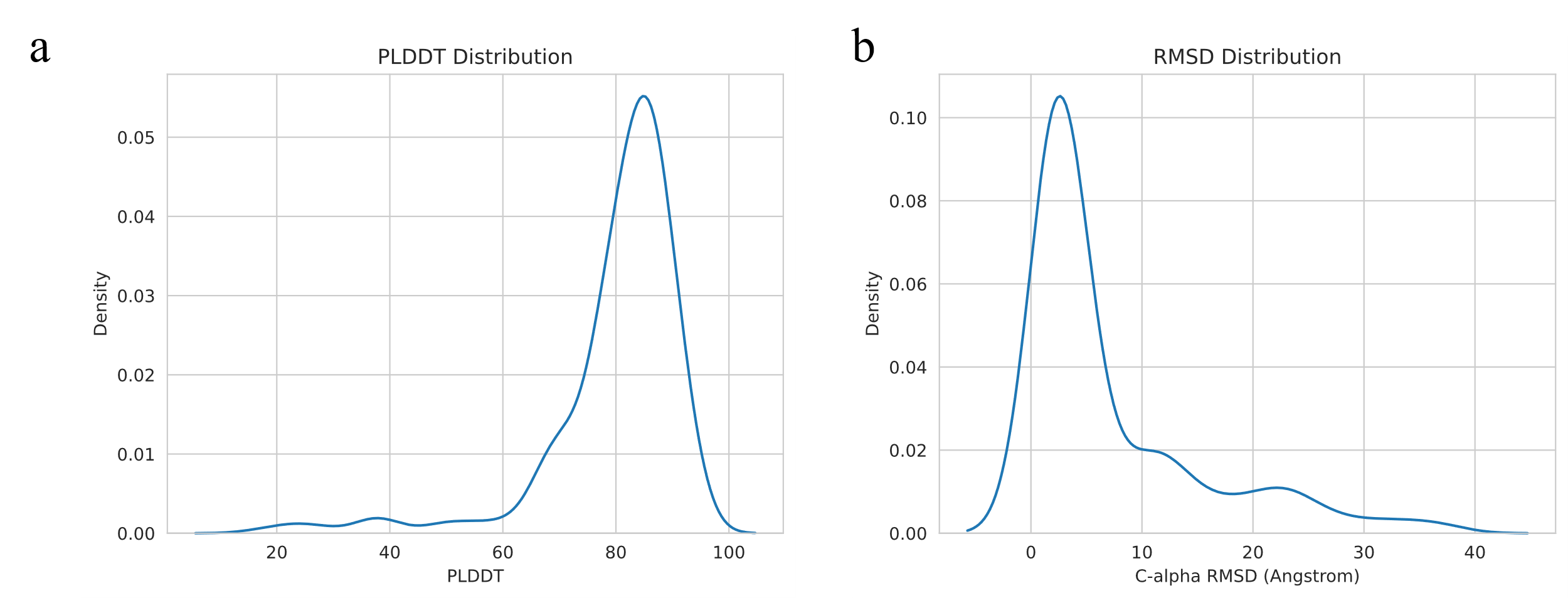
Fig.S4: PLDDT distribution and and C-α RMSD of predicted ESMFold structure. (a) PLDDT distribution. (b) Distribution of C-α RMSD between crystal structure and ESMFold predicted structure.

**S6. Failure examples of DUDE-AF2 test set**

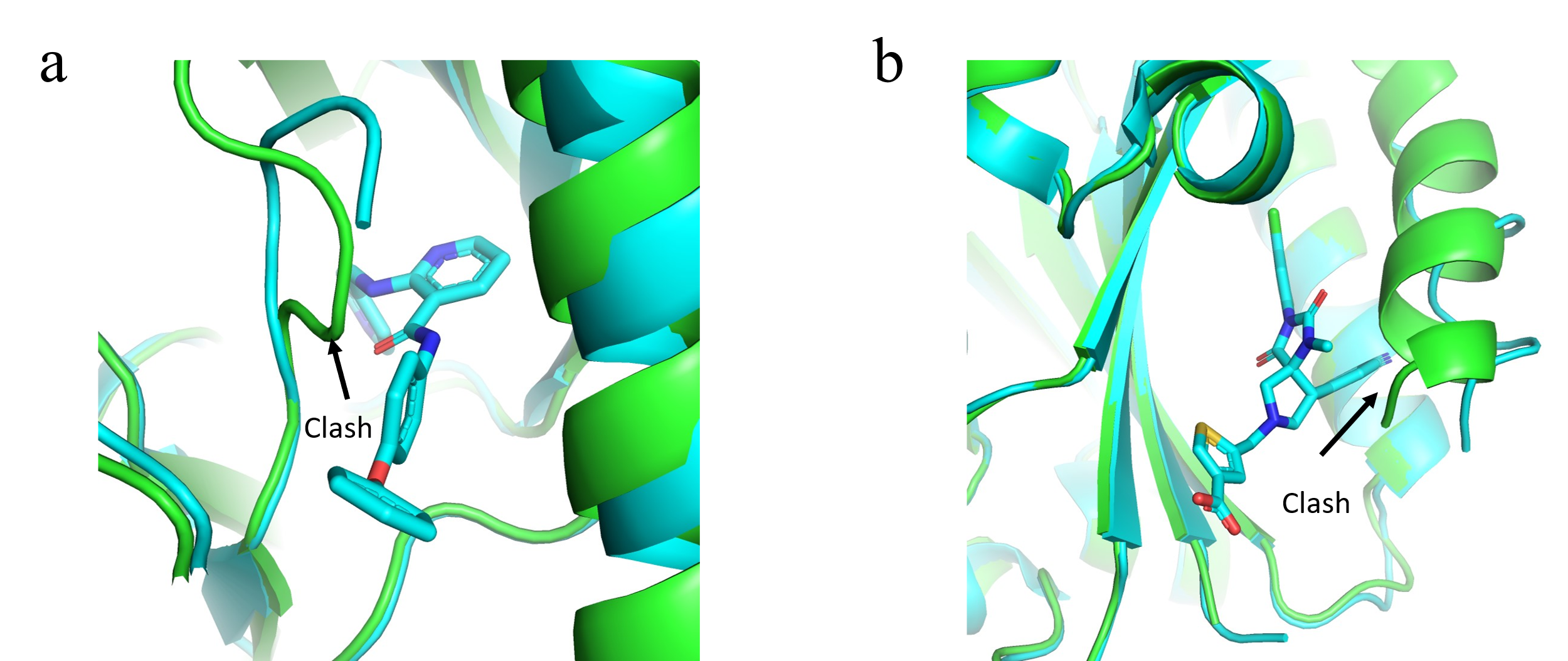

Fig.S5: Failure examples of target MET (a) and ITAL(b) in DUDE-AF2 test set. The cyan represent the crystal structure, the green represent the Alphafold2 modeled structure.
